# Optical Brain Pulse Monitoring for Detection of Large Vessel Occlusion Stroke in a Sheep Model

**DOI:** 10.1101/2025.01.30.635815

**Authors:** Jessica M. Sharkey, Elliot J. Teo, Sung W. Chung, Sally A. Grace, Jack Hellerstedt, Sigrid Petautschnig, Hiran A. Prag, Renée J. Turner, Thomas Krieg, Michael P. Murphy, Annabel Sorby-Adams, Barry Dixon

## Abstract

Early stroke detection and treatment are critical for improving patient outcomes. Optical brain pulse monitoring (OBPM) uses red and infrared light to capture brain pulse waveforms reflecting arteriole-to-venous pressure levels driving microvascular blood flow. This study assessed OBPM’s potential to detect middle cerebral artery occlusion (MCAo) and reperfusion in a clinically relevant sheep model. Stroke was induced in 11 Merino wethers via 4-hour occlusion of the right MCA, followed by 6 hours of reperfusion. OBPM recordings were taken at baseline, MCAo, early and late reperfusion. The OBPM brain pulse waveform classes were classified based on the presence of arterial or central venous circulation wave features. Magnetic resonance imaging assessed infarct volume at 2 hours post-reperfusion. Invasive brain tissue oxygen and intracranial pressures were also monitored. The OBPM brain pulse waveform classes changed during MCAo and reperfusion (*p* <0.0001). MCAo was associated brain pulses with venous circulation features (*p* = 0.0007). Reperfusion was associated with the return of arterial circulation features (*p* = 0.001). Early reperfusion was also associated with an increase in the brain pulse amplitude (*p* < 0.05) and the respiratory wave amplitude (*p* < 0.05). OBPM may aid in early stroke detection and reperfusion assessment following intervention.

## Introduction

The efficacy of stroke intervention and management relies on timely and accurate detection of evolving infarction.^1,2^ Existing interventions, including thrombolytics and endovascular thrombectomy (EVT), are dependent on timely administration, with every 60-minute delay in treatment resulting in death of an additional 120 million brain cells, worsening long-term outcomes.^3^

Current clinical guidelines recommend urgent brain computed tomography (CT) or magnetic resonance imaging (MRI) for patients with suspected stroke to determine stroke subtype and severity.^4,5^ Due to expense and complexity, these scanners are usually located in well resourced, metropolitan hospitals. Access is limited in rural, remote, and low-resourced environments, resulting in delayed stroke detection and treatment.^6^

A simple inexpensive point-of-care monitor could assist in earlier diagnosis and evaluation, and in conjunction with telemedicine stroke specialists, inform the decision for transport to a comprehensive stroke center for intervention. Furthermore, if large vessel occlusion (LVO) stroke was excluded, an informed decision about urgent patient transport may be avoided.

An approved simple point-of-care monitor to detect stroke does not currently exist, and alternative technologies including cerebral oximetry, electroencephalography (EEG) and transcranial ultrasound doppler have not demonstrated sufficient clinical efficacy to date.^7–10^ Mobile CT stroke units show promise, yet are only available in high population areas due to high running costs and capital expenditure, highlighting the need for more accessible technologies.

Optical brain pulse monitoring (OBPM) is a novel, non-invasive technique that uses red and infrared light to capture brain pulse waveforms, whose morphology reflects the relative arteriole to venous pressure levels that drive microvascular blood flow in the brain. The OBPM brain pulse classes include arterial expansion, arterial compression, hybrid, venous I and venous II, ranging from normal (arterial) to critically low blood flow (venous II).^11,12^ The ability of OBPM to observe these distinct pulse classes suggests potential to detect regional changes in cerebral blood flow following ischemic stroke. OBPM’s simplicity, affordability, and non-invasive nature thus make it an attractive technology for early stroke detection and monitoring with the potential for widespread use.

This study investigated the ability of OBPM to detect stroke and successful reperfusion in an ovine model. This model replicates key clinical features, including LVO via middle cerebral artery (MCA) occlusion and subsequent reperfusion. The large brain size of the sheep also facilitates concurrent non-invasive and invasive neuromonitoring.^13,14^ Using this model, we assessed OBPM signal changes during ischemic stroke and following reperfusion.

## Materials and Methods

### Institutional Approval

All procedures were approved by the Animal Ethics Committee of the South Australian Health and Medical Research Institute (SAM-21-098). Experiments complied with the Australian NHMRC Code for the Care and Use of Animals for Scientific Purpose (8^th^ edition, 2013) and were reported according to the Animal Research Reporting of In Vivo Experiments (ARRIVE 2.0) guidelines.^15^

### Experimental design

Eleven Merino wethers (*Ovis aries,* 36-42 months, 52.0-69.5 kg) underwent four-hour right MCA occlusion followed by a reperfusion period of 6 hours (see Figure 1A). The OBPM sensor was placed ipsilateral to the MCA occlusion in all animals. The contralateral hemisphere was also monitored in six animals. Invasive brain tissue oxygen monitoring (PbtO_2_) was monitored in the ipsilateral hemisphere, while invasive intracranial pressure (ICP) was monitored in the contralateral hemisphere (Figure 1B).

**Figure 1.**
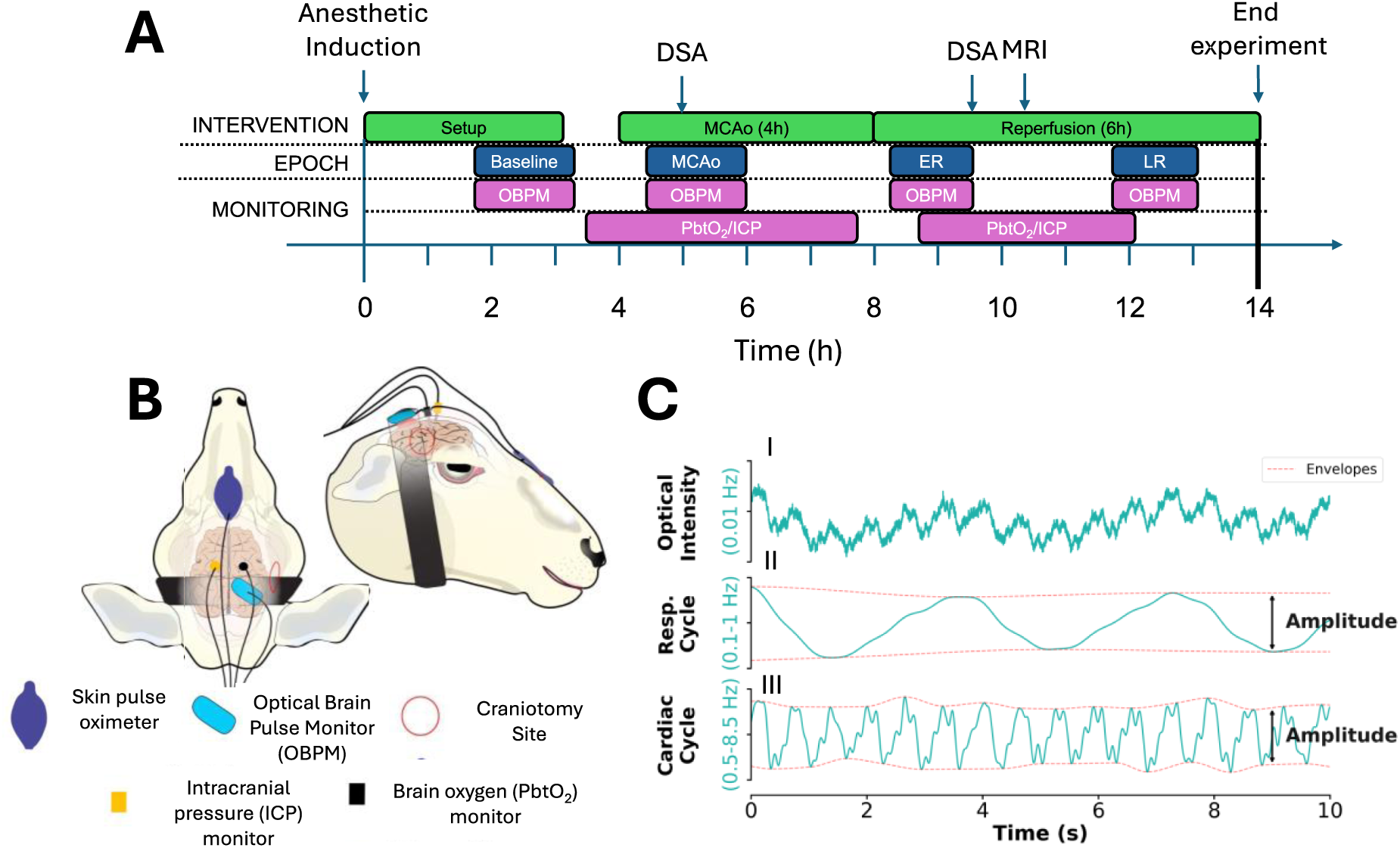
Experimental set up and optical brain pulse monitor outputs. (**A**) Timeline illustrating the sequence of events, and the periodic monitoring over the course of the experimental protocol. (**B**) The illustration shows the placement of monitoring devices on the sheep’s head. The craniotomy site is marked in red, the skin pulse oximeter and optical brain pulse monitor (OBPM) in blue and light blue respectively, the invasive brain oxygen monitor in black, and the intracranial pressure monitor in orange. (**C**) Illustration of the decomposition of the (**I**) raw OBPM waveform into (**II**) respiratory and (**III**) cardiac components. The red line illustrates the signal envelope from which the amplitude is derived. *Abbreviations: OBPM, optical brain pulse monitor; ER, early reperfusion; LR, late reperfusion ICP, invasive intracranial pressure; MRI, magnetic resonance imaging; MCAo, middle cerebral artery occlusion; PbtO_2_, invasive brain tissue oxygen; s, seconds; h, hours*.

OBPM recordings were undertaken at four timepoints: baseline (prior to neurosurgical procedures), during MCA occlusion (∼30 minutes after onset), early reperfusion (∼30 minutes post-reperfusion), and late reperfusion (∼4 hours post-reperfusion), each for approximately 30 minutes. Animals were sacrificed under isoflurane anesthesia via perfusion exsanguination at six hours post-reperfusion.

### Anesthesia and physiological monitoring

Anesthesia was induced with ketamine (0.05 mL/kg, 100 mg/kg, CEVA, Australia) and diazepam (0.08 mL/kg, 5 mg/mL, Pamlin, CEVA, Australia). Animals were intubated and positioned supine. An arterial catheter was placed in the right common carotid artery for blood pressure monitoring and blood gas sampling, and a catheter in the left jugular vein for fluid and drug administration. Anesthesia was maintained with isoflurane (1.5-2.0%, Henry Shein, Australia) and ketamine (4 mg/kg/hr, CEVA, Australia). Administration of crystalloid fluids (Baxter Health, Australia) via jugular line and sodium chloride (Baxter Health, Australia) via carotid line ensured hydration and catheter patency, respectively. Arterial blood samples were collected hourly and analyzed (OPTI-3 CCA Electrolyte and Blood Gas Analyzer, Roche, USA). Esophageal temperature, end-tidal CO_2_, pulse rate, and mean arterial blood pressure were continuously monitored.

### Surgical approach

The ovine model of MCA occlusion (MCAo) was implemented in this study as previously described (for full details see Supplementary Detailed Method 1).^13,16,17^ Briefly, animals were positioned prone for a right MCA approach via pterional craniotomy with a high-speed pneumatic drill (Midas Rex, Medtronic, USA). Following durotomy, a temporary aneurysm clip (Aesculap YASARGIL Aneurysm Clip, Germany) was applied to the proximal M1 branch of the MCA (Figure 2A). Synthetic dural regeneration matrix (Durepair, Medtronic, Australia) was applied over the exposed pial surface, autologous bone repositioned, and muscles reopposed. Reperfusion was achieved after four hours by re-opening the surgical site and removing the aneurysm clip. Successful reperfusion was confirmed with DSA (Figure 2B). A permanent cranioplasty was performed thereafter with autologous bone and polyacrylic cement (PMMA, Lang Dental, USA).

**Figure 2.**
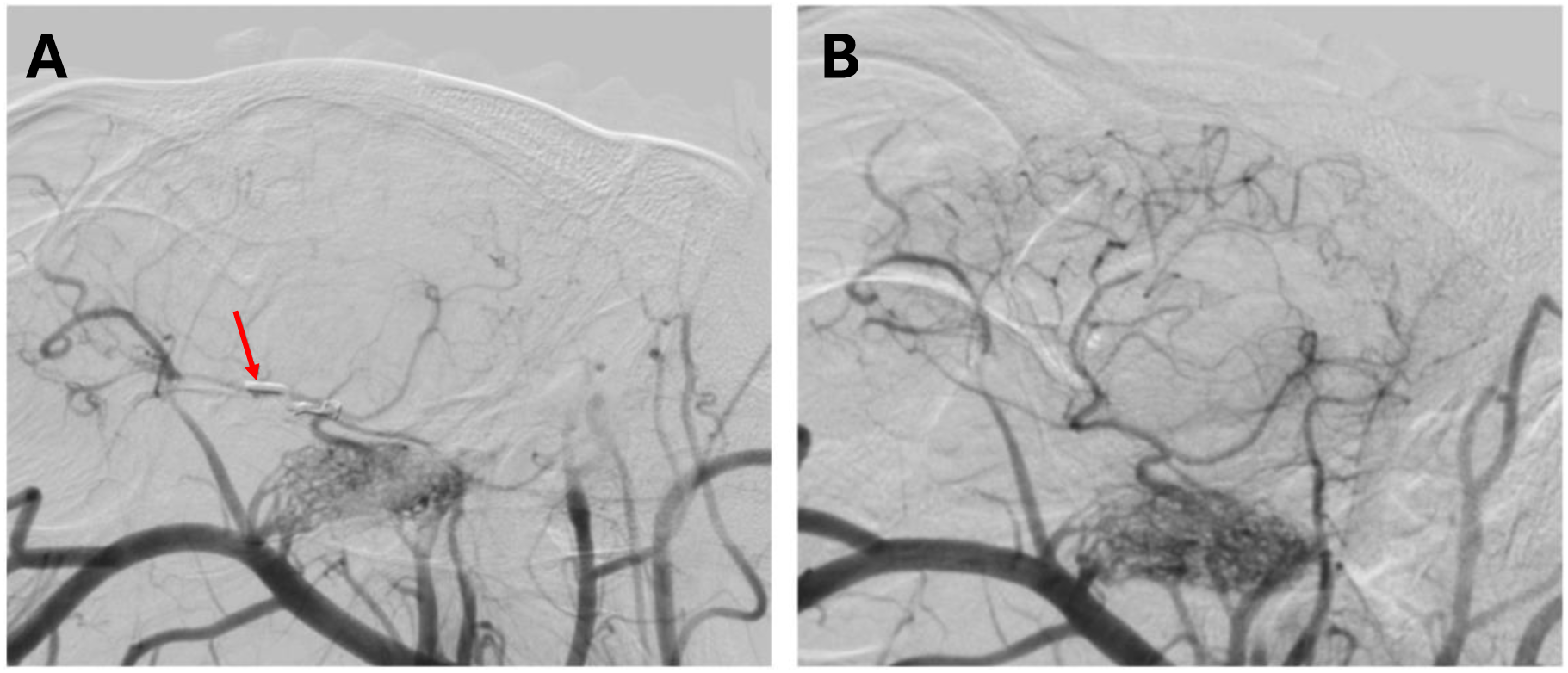
Middle cerebral artery occlusion (MCAo) and reperfusion on cerebral digital subtraction angiography. **A)** Occlusion of the proximal M1 segment of the MCA was confirmed following aneurysm clip application (red arrow), resulting in reduced blood flow to the MCA territory. **B)** Reperfusion was confirmed by repeat angiography following clip removal. *Abbreviations: middle cerebral artery, MCA*.

### Neuroimaging

One hour following MCAo, animals were transported to a catheter laboratory for digital subtraction angiography (DSA) to confirm successful MCA occlusion (Siemens AXIOM-Artis, Germany). Following 1 hour of reperfusion, DSA was reacquired to confirm vascular reperfusion and MCA patency, after which animals were transported to an MRI suite for neuroimaging (3 T Magnetom Skyra, Siemens, Germany) under anesthesia (3% isoflurane). Diffusion-weighted imaging (DWI; *b*1000 s/mm²) was acquired to evaluate infarct volume with an echo time of 1 ms, repetition time of 5790 ms, acquisition matrix of 190 × 190, field-of-view of 163 × 163 mm, and slice thickness of 3 mm. Infarct volume was calculated using semi-automated three-dimensional segmentation tools in ITK-SNAP v4.0.2.^18,19^

### Optical Brain Pulse Monitoring

The OBPM has been previously described in detail.^2,11,12,20,21^ Briefly, the device employs red (660 nm) and infrared (940 nm) light sources, selected for their differential absorption to oxyhemoglobin and deoxyhemoglobin. Comparable to traditional skin-based pulse oximetry, oxyhemoglobin absorbs more infrared, whereas deoxyhemoglobin absorbs more red light. The sensor consists of a light-emitting diode and a photodetector. The geometry of the sensor preferentially captures photons reflected from the pial venules that lie on the cortical surface of the brain.^11^

The OBPM brain pulse shape reflects pulsatile blood volume changes in pial venules that lie on the cortical surface within the sub-arachnoid space (Figure 3). The pial venules blood volume is 4-fold higher than in the capillary beds of the cortex.^22^ The pial venules act as a blood reservoir.^23–25^ Thus, a very large venous blood volume lays on the surface of the cortex, providing a strong OBPM optical signal.

**Figure 3.**
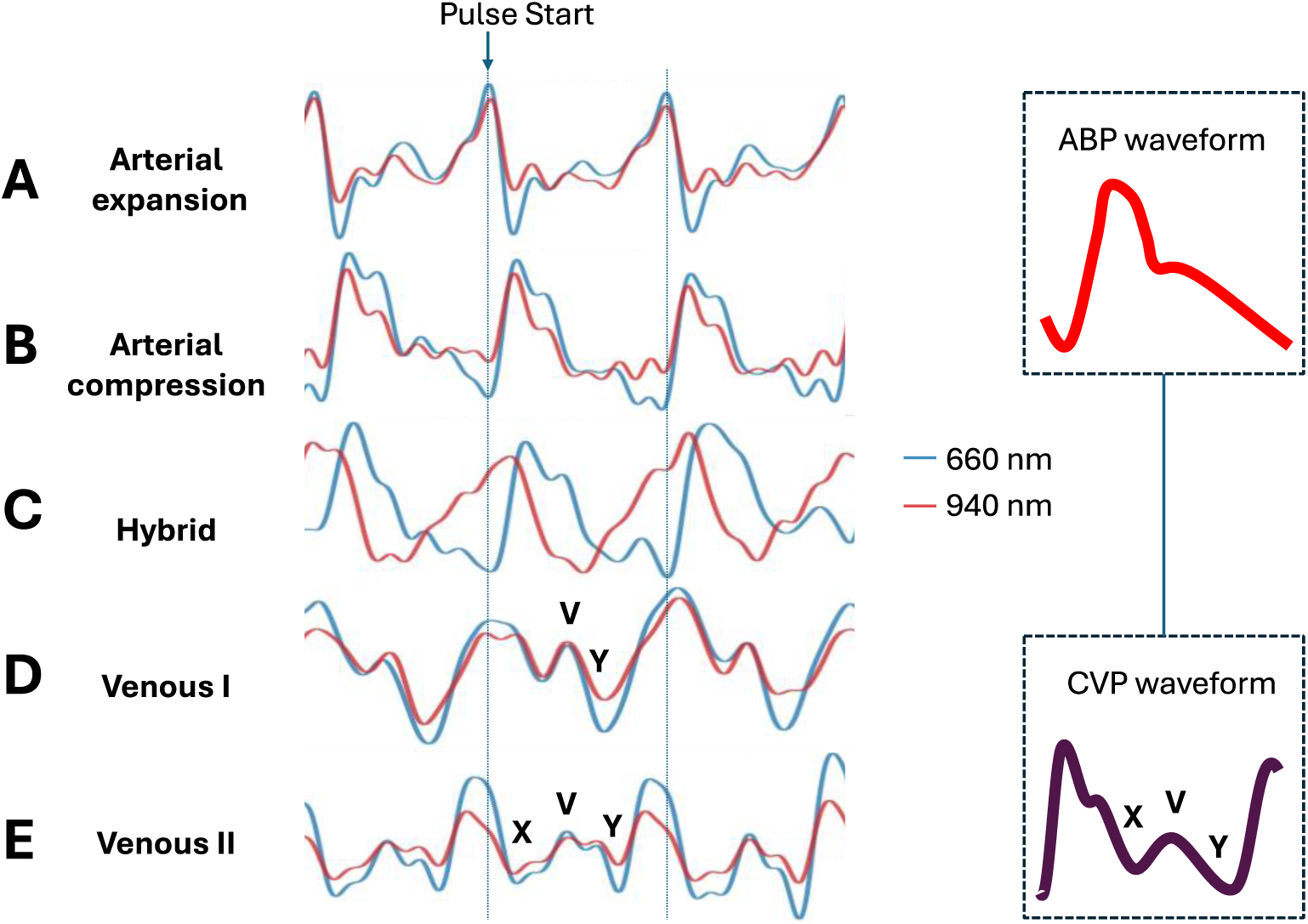
Morphology of OBPM brain pulse classes. (A) arterial expansion, (B) arterial compression, (C) hybrid, (D) venous I, and (E) venous II brain pulses and comparison to arterial blood pressure (ABP) and cerebral venous pressure (CVP)_ waveforms. Red brain pulse is 940 nm and blue brain pulse 660 nm. *Abbreviations: ABP, arterial blood pressure; CVP, central venous pressure*.

For all animals, the OBPM sensor was placed on the head overlying the right MCA territory. The position was indicated with surgical marker following the baseline reading for consistent replacement for subsequent recordings, as the sensor was removed between data collection periods. An additional reflectance pulse oximeter (Nellcor ^TM^, Medtronic, USA) was placed on the midline over the frontal sinuses to provide reference photoplethysmogram, respiration, and pulse rates.

### PbtO_2_ and ICP monitoring

Bilateral 5 mm burr holes were made in the parietal bones using a hand drill (Raumedic, Germany) for ipsilateral PbtO_2_ (Neurovent-TO probe, Raumedic, Germany) and contralateral ICP (Neurovent-P probe, Raumedic, Germany) probe insertion. Data were recorded using the Raumedic MPR2 logO system (Raumedic, Germany). A two-point calibration (0 and 100 mmHg) was performed before inserting the probes and at the end of the recording period to confirm measurement accuracy. Due to lack of MRI compatibility, probes were removed and re-inserted at the original depth before and after MRI acquisition.

### OBPM, ICP and PbtO_2_ signal processing

OBPM data was processed using SciPy v1.10^26,27^ and proprietary scripts deployed in Python v3.10 (for full details see Supplementary Detailed Method 2). Raw data were cleaned using an automated pipeline to remove artifacts such as photodetector saturation, temporary disconnection, and motion. The clean data were filtered to highlight specific frequency components relevant to different physiological mechanisms for quantitative analyses. For PbtO_2_ and ICP recordings, sections of data were rejected based on manual screening, where the criteria for rejection were similar to those for automatic filtration of the OBPM signal. Filtered skin photoplethysmogram was used as an aid to identify the onset of the cardiac pulse. Data was captured at a rate of 500 Hz and stored for offline processing.

### Qualitative OBPM waveform assessment

To assess the OBPM waveform classes, all recordings were examined by three expert observers blinded to both the animal number and timepoint. Conflicts were resolved by consensus agreement. The OBPM brain pulse waveform classes were assessed based on the presence of arterial or central venous pressure wave features.^11^

Under physiological conditions, the brain pulse shape resembles an arterial pressure wave. There are two classes of arterial brain pulses—the arterial expansion pulse and the arterial compression pulse.^28–34^ The terms expansion and compression refer to the likely blood volume changes in the pial venules during the systolic phase of the cardiac cycle. The arterial expansion pulse (Figure 3A) has a more prominent arterial character than the compression. This pulse may result from pial venules with relatively high blood pressure that expand during systole resisting compression from the simultaneously expanding brain cortex. This gives rise to a fall in light intensity during systole.^35,36^ The arterial compression pulse (Figure 3B) is essentially a mirror image of the expansion pulse, however the light intensity increases during early systole. This waveform likely reflects pial venules with low blood pressure that are compressed during systole from the aforementioned expanding brain cortex.^28–34^

Brain pulse classes associated with low microvascular blood flow include hybrid, venous I and venous II pulses (Figure 3C). The hybrid pulse likely represents a state in which the cortical arteriole pressure is low yet still exceeds venule pressure. This pulse waveform is characterized by distinct shapes for the 940 nm and 660 nm wavelengths, due to their different responses to low blood oxygen levels. The venous I pulse (Figure 3D) likely reflects a state where arteriole and venous pressure levels are similar which gives rise to a pulse with somewhat undifferentiated features during systole. This pulse may demonstrate more obvious central venous pressure features in the diastolic phase including A and Y waves, as arteriole pressure levels fall during diastole. The venous I pulse is consistent with very low microvascular blood flow and potentially no flow during diastole. The venous II pulse (Figure 3E) has clear central venous pressure features such as A, C, V, X and Y waves, and is consistent with venous pressure exceeding arteriole pressure throughout the entire cardiac cycle.^11^

### Quantitative OBPM assessment

To evaluate the relationship between evolving pathology and OBPM signal frequency components we analyzed the amplitude of the cardiac oscillation, the amplitude of the respiratory oscillation, and the unfiltered optical intensity at each timepoint (Figure 1C). The OBPM respiratory signal results from dynamic changes in brain cerebrospinal fluid (CSF), venous and arterial blood flow, driven by thoracic and abdominal pressure oscillations during respiration. Slower frequency oscillations (0.5–3 waves/minute) that occur in brain injury states such as slow waves (Lundberg B waves) due to oscillations in cerebral blood flow were also assessed, as were changes in the optical intensity (the raw unfiltered optical signal).^21,37^

### Statistical analysis

Analyses were performed in Python v3.10, R Studio v0.84.1, and GraphPad Prism v10.4.1. Interrater reliability of waveform classes was performed between all assessors and reported using the Fleiss κ statistic. Assessment of the brain pulse class responses to MCAo and reperfusion were converted to an ordinal scale, where 5 = arterial expansion, 4 = arterial compression, 3 = hybrid, 2 = venous I and 1= venous II. Repeated measures one-way analysis of variance (ANOVA) assessed the change in the mean values for each timepoint, where sphericity was not assumed. Sidak’s multiple comparisons test was used to compare means between timepoints. A Generalized Estimating Equation (GEE) was used to model changes in OBPM signal components, including the cardiac amplitude, the respiratory amplitude, optical intensity, PbtO_2_ and ICP. Additional pairwise comparisons were made between MCAo and both early and late reperfusion. The Benjamini-Hochberg method was used to control the false discovery rate. Statistical significance was accepted at *p* < 0.05.

## Results

### Infarct volume, PbtO_2_ and ICP

All animals were included in the final analysis, except where explicitly stated. Physiological variables were maintained within normal ovine limits (Supplementary Table 1). At ∼ 2 hours following ischemia the median DWI infarct volume was 8.3 cm^3^ (interquartile range 5 - 28.9 cm^3^).

Complete PbtO_2_ data was available in 7 sheep. A significant decrease in the mean ipsilateral PbtO_2_ (*p* < 0.001) was observed following MCAo compared with baseline (35 to 8 mmHg). Levels increased to 28 mmHg following early reperfusion (*p* < 0.01, compared with MCAo) and increased to 38 mmHg by late reperfusion (Figure 4). Complete ICP data were available in 10 sheep. A significant decrease in the mean contralateral ICP (*p* = 0.01) was observed following MCAo compared with baseline (11 to 7 mmHg). Levels were 4 mmHg during early reperfusion and 7 mmHg by late reperfusion (Figure 3).

**Figure 4.**
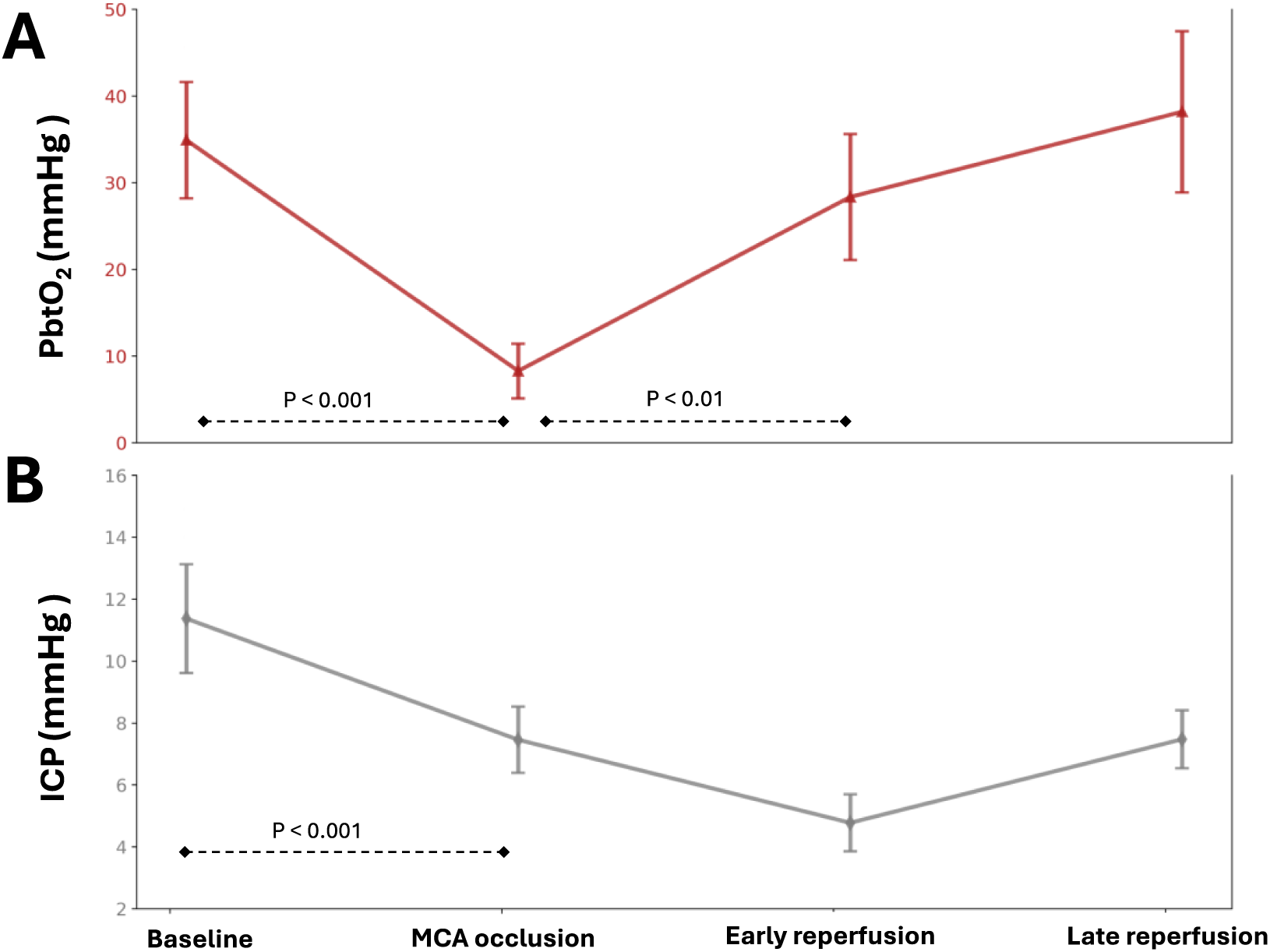
PbtO_2_ and ICP measurements throughout the experiment. **A)** PbtO_2_ levels in the ipsilateral stroke hemisphere fell significantly during MCAo, increased during early reperfusion, and returned to baseline levels by late reperfusion. **B)** ICP levels in the contralateral hemisphere fell significantly during MCAo. Data are presented as mean ± SE. *Abbreviations: PbtO_2_ partial pressure of brain oxygen; ICP, intracranial pressure; MCAo, middle cerebral artery occlusion; mmHg, millimeters of mercury*.

### OBPM waveform class responses to MCA occlusion and reperfusion

Classification of the predominant waveform class at each timepoint demonstrated moderate agreement between assessors (κ = 0.433, *p* < 0.0001). Ipsilateral OBPM brain pulse classes were assessed in all 11 animals and changed during both MCAo and reperfusion timepoints (*p* < 0.0001, Figure 5). At baseline the brain pulse classes were predominately arterial in character (73% of sheep). The classes were arterial expansion (36%), arterial compression (36%), hybrid (9%), and venous I (18%). During MCAo, the brain pulse classes developed venous circulation features in 64% of sheep (*p* = 0.0007, compared with baseline). The classes were arterial expansion (18%), arterial compression (9%), hybrid (9%), venous I (36%) and venous II (27%). During early reperfusion, the brain pulse classes transitioned to predominately arterial circulation features in 91% of sheep (*p* = 0.001, compared with MCAo). The classes were arterial expansion (55%), arterial compression (36%), and venous I (9%). At late reperfusion, the predominate brain pulse classes remained arterial (91% of sheep), and included arterial expansion (73%), arterial compression (18%), and hybrid (9%). The venous II class was only present during MCAo. Contralateral OBPM classes were assessed in six sheep. No significant change in the contralateral brain pulse classes were observed between any of the timepoints (all p > 0.05; Supplementary Figure 1).

**Figure 5.**
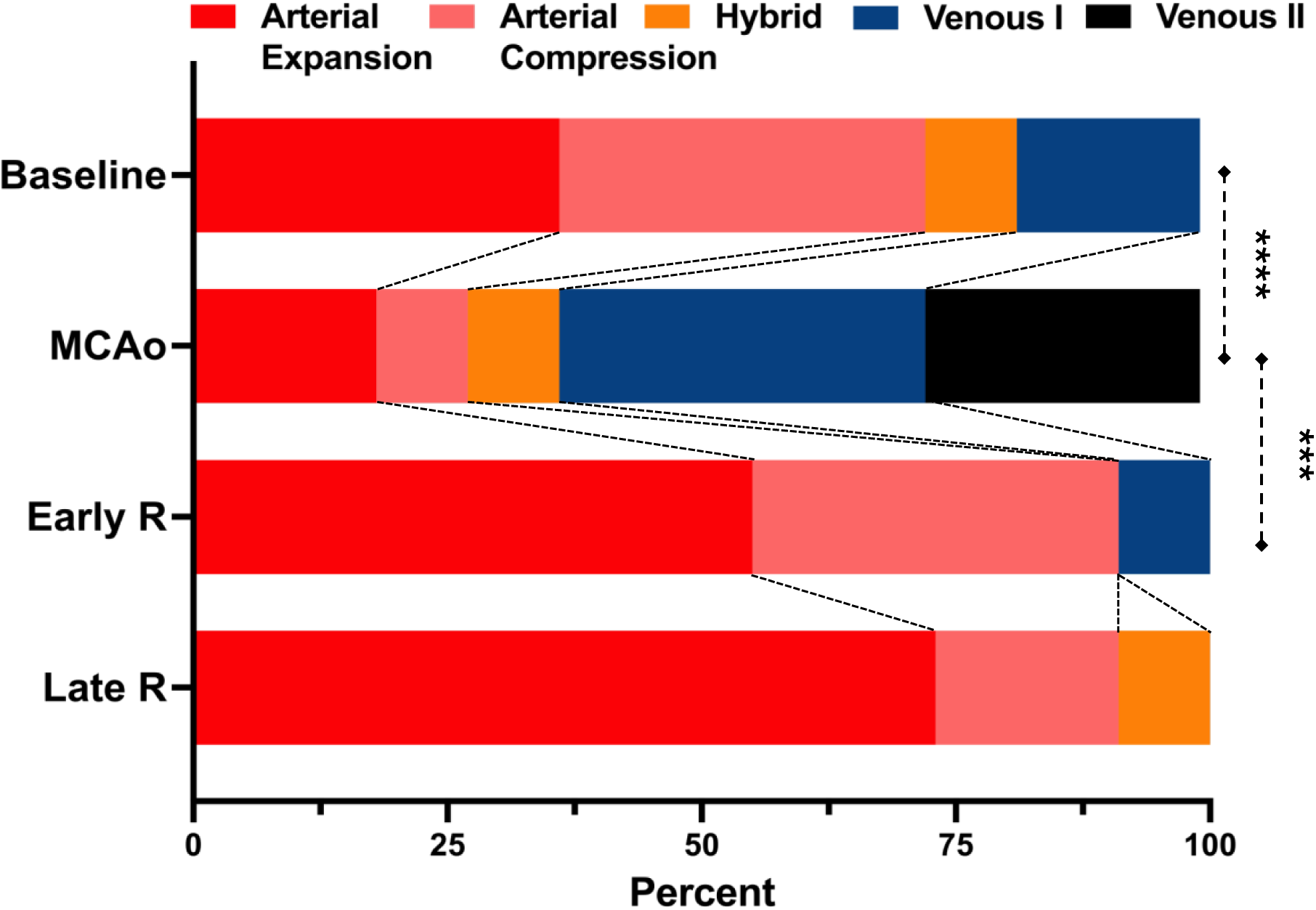
Brain pulse waveform classes change in response to MCAo and reperfusion. During the MCAo the brain pulse waveforms developed predominately venous circulation features (Venous I and Venous II brain pulse classes). At early reperfusion, the brain pulse waveforms transitioned back to predominately arterial brain pulse classes. *Abbreviations: MCAo, middle cerebral artery occlusion; Early R, early reperfusion; Late R, late reperfusion. *** p* = 0.001, **** *p* = 0.0007.

### Optical Intensity

The optical intensity was higher for both the red and infrared light sources during early reperfusion compared with MCAo (*p* < 0.05, Figure 6A). The amplitudes of the OBPM cardiac pulse and the respiratory waves increased following early reperfusion compared with MCAo for both the red and infrared light sources (*p* < 0.05, Figure 6). In two animals the respiratory waves in the ipsilateral and contralateral hemispheres were out of phase following reperfusion and delayed over the ipsilateral hemisphere (Figure 7A).

**Figure 6:**
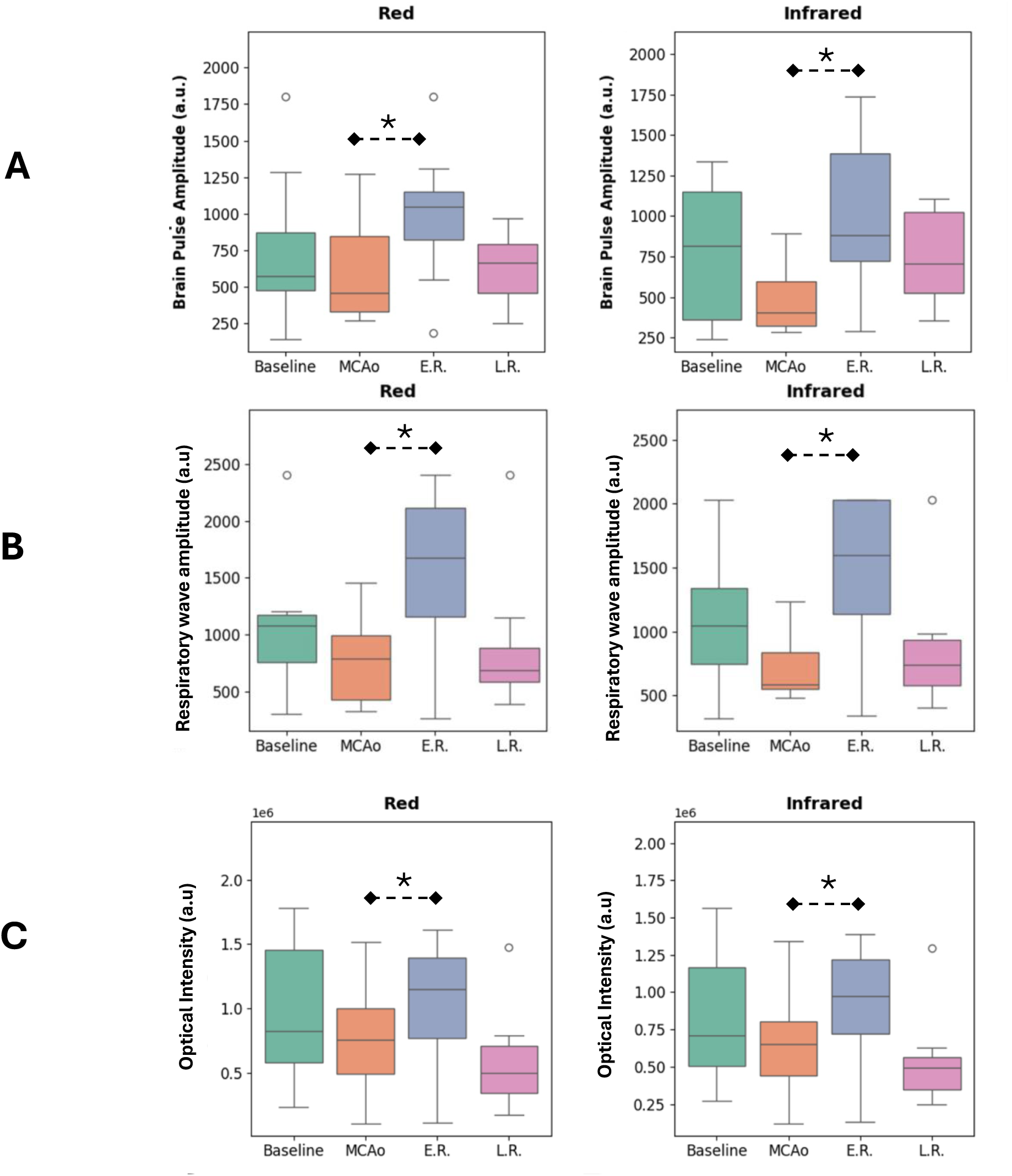
OBPM output changes in response to MCAo and reperfusion. Box plot of changes in A) cardiac pulse amplitude, B) respiratory wave amplitude and C) optical intensity in the stroke hemisphere for the red (660nm) and Infrared (940 nm) wavelengths. A significant increase in amplitudes were observed between MCAo and early reperfusion for each of these signal outputs. The box-and-whisker plot shows the median, interquartile range (box), and spread (whiskers), with outliers as individual points. *Abbreviations: a.u., arbitrary units; MCAo, middle cerebral artery occlusion; E.R. early reperfusion; L.R., late reperfusion.* * *p <* 0.05).

**Figure 7:**
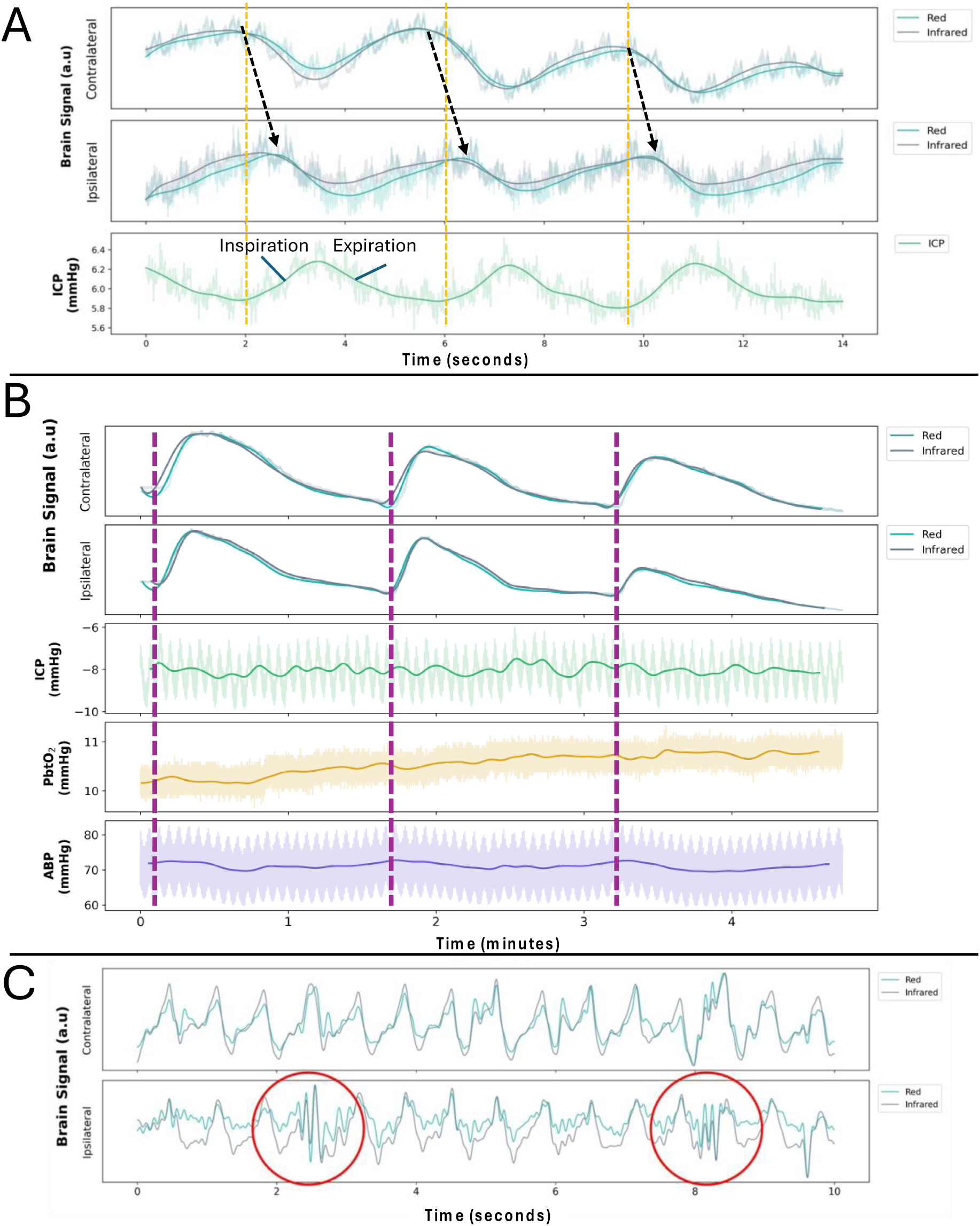
Illustration of non-cardiac OBPM signal features. **(A)** OBPM respiratory wave phase asynchrony between ipsilateral stroke and contralateral hemispheres. The stroke hemisphere respiratory wave was delayed relative to the contralateral hemisphere (black arrows). The oscillations in ICP indicate the respiratory phases (orange lines). The OBPM and ICP signals are faded for visualization purposes, a filter highlighting the respiratory osscilation has been applied. **(B)** Bilateral slow waves during early reperfusion. Slow-wave pattern oscillations were observed in both the ipsilateral and contralateral hemispheres following reperfusion, with a slow wave frequency of approximately 1 per minute. Local increases in ABP were associated with slow wave troughs (magenta lines) but had no clear association with ICP or PbtO_2_. **(C)** Episodes of fast waves (spindles) during late reperfusion. The red circles highlight fast waves in the stroke hemisphere, with an oscillation frequency of ∼12 Hz, occurring sporadically every 10-45 seconds. These may reflect fast oscillations of cerebrovascular smooth muscle with transmitted pressure waves as a possible mechanism. *Abbreviations: ABP, arterial blood pressure; a.u., arbitrary units; ICP, intracranial pressure; mmHg, millimeters of mercury; min, minutes; OBPM, optical brain pulse monitoring; PbtO2, partial pressure of brain tissue oxygen; s, seconds. Infrared light: 940 nm; Red light: 660 nm*.

### Slow waves

When considering the waveforms from individual animals, one sheep demonstrated Lundberg B/slow waves in the stroke hemisphere ∼30 minutes following MCAo, though contralateral OBPM was not performed for comparison. Five sheep developed slow waves during early reperfusion, and of these, 4 demonstrated persistent slow waves out to late reperfusion. These slow waves were often, but not always associated with modulations in invasive ICP and PbtO_2_. During the slow wave episodes, two sheep exhibited loss of the brain pulse at the peak optical intensity which returned as the optical intensity fell to baseline.

### Fast waves

One animal demonstrated an unusual waveform feature resembling an electroencephalographic spindle like activity in the ipsilateral hemisphere during late reperfusion. The oscillation frequency was fast at approximately 12 Hz, occurring sporadically every 10-45 seconds. The spindle like activity could not be attributed to facial movement, sensor instability or any other externally driven artefact (Figure 7C).

## Discussion

Our study demonstrates that non-invasive OBPM can detect hemodynamic changes associated with ischemic stroke and subsequent reperfusion in an ovine model. During MCAo, the OBPM brain pulse shape developed venous circulation features. Following successful reperfusion, the OBPM brain pulse recovered arterial circulation features resembling those seen at baseline. Our findings suggest this novel monitor has potential as a point-of-care device for early stroke detection, with the ability to detect successful reperfusion following intervention.

### The OPBM waveform developed venous circulation features during MCA occlusion

We believe the major cerebrovascular factor influencing the OBPM waveform shape is the relative arteriole to venous pressure across the pial venule microvascular bed, driving microvascular blood flow. During MCAo arteriole pressure drops below venous pressure levels.^38^ In this setting the brain pulse waveform may develop venous circulation features as the venous pressure becomes the higher pressure oscillating the pial venous blood.^11^

During MCAo, we found the venous I and venous II pulse classes predominated. The venous II pulse features were consistent with venous pressure exceeding arteriole pressure throughout the cardiac cycle which may result in retrograde or reversal of venous blood flow. Reversal of blood flow in cerebral veins has been demonstrated in critically ill patients with severe brain injury using transcranial doppler;^39–41^ in cortical arteriole capillary beds and venules in rat models of vascular injury and stroke;^42–45^ and in a human cadaver study in response to raised jugular venous pressure.^46^ Notably, pooling of blood in the ipsilateral cerebral veins is a feature of acute stroke and is associated with poor patient outcomes.^47,48^ Pooling could reflect filling from the extensive communications of the venous circulation within and between hemispheres to the very low venous pressure vessels within the ischemic territory.^49,50^

In a first-in-human clinical study in 11 patients presenting with LVO stroke, we also found venous circulation features were present in the brain pulse waveforms.^51^ Similarly, in a study of critically ill brain injured patients following prolonged out of hospital cardiac arrest we observed extended periods of venous II brain pulses which were associated with severe brain injury and poor neurological outcomes.^11^ These findings suggest that OBPM could assist in earlier diagnosis and treatment of LVO stroke.

### The OPBM waveform arterial circulation features returned following reperfusion

PbtO_2_ recovered to baseline levels following reperfusion indicating successful recovery of blood flow to the ischemic territory as observed on DSA. The OBPM waveform also changed following reperfusion, regaining arterial circulation features. The proportion of sheep with the arterial expansion class increased from 36% at baseline to 55% during early reperfusion and further to 73% during late reperfusion. Animal models of stroke found reperfusion maybe associated with hyper-perfusion which could increase pial venous blood pressure. This could be a factor in the high proportion of sheep developing the arterial brain expansion waveform class following reperfusion.^35,38,52^

Other OBPM signals also changed with reperfusion, including increases in the brain pulse amplitude and respiratory wave amplitude. The increase in brain pulse amplitude could reflect greater pulsatile blood flow due to the restored perfusion pressure and vessel compliance, which enhance the transmission of cardiac-driven blood volume changes. ^11,38^ Similarly, the increase in respiratory wave amplitude arises from the enhanced transmission of thoracic pressure oscillations through the cerebral vasculature, which becomes more pronounced with elevated cerebral blood flow and increased cortical blood volume.^38^

The OBPM optical intensity from the stroke hemisphere for both the red and infrared light sources also increased during early reperfusion. This was surprising as, discussed above, blood volume would be expected to increase, and more blood would absorb more light leading to a reduction in the optical intensity signal. The blood volumes in the brain are however compartmentalized. We believe the blood volume in the outer pial venules is the predominate signal source of the OBPM signal. Thus, it is plausible that this volume might decrease as a result of increased cerebral perfusion with a net reduction in venous blood pooling.^47,48^

Our findings suggest that OPBM could also provide a method to monitor microvascular blood flow responses to thrombolytic or EVT intervention. The technical success of EVT is currently measured by the achievement of reperfusion, evaluated using the Modified Treatment in Cerebral Ischemia (mTICI) scale which is qualitatively estimated by the interventionalist at the end of the procedure using cerebral DSA. However, there remains a gap between successful arterial reperfusion as assessed by mTICI grade and patient outcomes, as almost 50% of patients do not experience favorable outcomes despite successful thrombectomy.^53–56^ The mTICI score has limitations in assessing microvasculature reperfusion which may be an important factor in the discrepancy between a good mTICI grade yet poor patient outcome. OBPM could bridge this gap by providing real-time, quantitative assessments of microvascular perfusion during and after EVT, enabling clinicians to directly measure the success of reperfusion at the microvascular level and potentially improving patient outcomes.

### Respiratory Oscillations

In two animals, we found that the respiratory waves in the ipsilateral hemisphere were delayed compared to the contralateral hemisphere. Normal respiratory waves monitored by OBPM are similar in phase and amplitude over both hemispheres^11^, with the respiratory waves resulting from brain movement due to oscillations in CSF and venous drainage during the respiratory cycle.^57–64^ MRI studies in traumatic brain injury demonstrate large unilateral hemispheric respiratory movements of the brain.^59,65^ Infarction with unilateral cerebral edema could give rise to abnormal respiratory motions as we have previously observed in critically ill patients with severe brain injury.^11^

### Slow Waves

Slow waves commonly developed following reperfusion. These are likely to represent Lundberg B waves which have a frequency of 0.5–3 waves/min.^66^ Lundberg B waves are seen in a range of brain injuries such as subarachnoid hemorrhage, traumatic brain injury, and ischemic stroke.^67^ Studies using transcranial doppler ultrasound, found Lundberg B waves arise from brief periods of increased MCA blood flow due to vasodilation, which might serve a dual purpose: facilitating glymphatic transport of waste products and supporting tissue repair by increasing local perfusion.^68,69^ These mechanisms underscore the potential importance of Lundberg B waves in the recovery and maintenance of brain homeostasis after injury.

### Fast Oscillations

The brain is a soft but muscular organ with smooth muscle present on all cerebral arteries, arterioles, capillaries (pericytes), venules and veins.^70,71^ We speculate that spasms (fast oscillations) of cerebrovascular smooth muscle could be a mechanism of the fast waves we observed in one animal. These fast waves appeared similar in character to Spindles a seen in EEG recordings. These fast waves may influence cerebral autoregulation and contribute to a cascade of secondary injury processes, particularly in regions already vulnerable due to primary injury or edema.^11,28,72,73^ Future studies should aim to elucidate the genesis and consequences of these oscillations, as well as their potential role in the progression or resolution of brain injury.

### Limitations

This study has several limitations that warrant consideration. The small sample size limits the generalizability of our findings and underscores the need for larger-scale studies. Additionally, the classification of brain pulse waveforms, while performed by blinded investigators, involves inherent subjectivity due to the continuous nature of waveform changes. This subjectivity introduces a potential for misclassification, which could be mitigated in future studies by objective signal processing and classification techniques.

The positioning of the sensor on the sheep skull was constrained by the need to avoid regions near the horn buds, where the skull is significantly thicker, particularly in male animals. As a result, the sensor placement may not have always directly overlaid the MCA territory. This limitation is compounded by differences between the sheep model and human stroke, as sheep lack the pre-existing cerebrovascular conditions often seen in humans, such as poor collateral blood flow. The sheep model also required craniectomy to apply the aneurysm clip to the MCA and cannulation of the carotid artery which could cause iatrogenic cerebral and extracranial tissue injury.

We found that at baseline several sheep demonstrated brain pulse classes associated with low cerebral blood flow. It is unclear if this represents a normal state in the sheep or if it is associated with the surgical approach, such as a reduction in vascular resistance due to inhalational isoflurane anesthesia. Finally, the OBPM sensor was removed between monitoring time points for logistical reasons, introducing some potential intra-individual variability in the brain region monitored.

### Conclusions

This study highlights the potential of OBPM as a novel tool for real-time stroke detection and monitoring. The OBPM brain pulse develops venous circulation waveform features during MCAo. Following reperfusion arterial circulation waveform features return. OBPM may be useful as a point of care device for early LVO stroke detection and could also assist in assessment of reperfusion following stroke treatment with thrombolytics or EVT.

This preliminary, preclinical study supports the utility of OBPM in stroke management and suggests its broader potential in monitoring cerebrovascular changes in neurological conditions. Future work should validate these findings in human populations and refine the technology for clinical use.

## Author Contributions

J.M.S: Conceptualization, Data curation, Investigation, Methodology, Resources, Visualization, Writing – review & editing.

E.J.T: Formal analysis, Visualization, Data curation, Writing – original draft, Writing – review & editing.

S.W.C: Formal analysis, Visualization, Data curation, Software, Writing – review & editing. S.A.G: Formal analysis, Software.

J.H: Software, Writing - review & editing.

S.P: Data Curation, Formal analysis, Writing - review & editing.

H.P: Funding acquisition, Investigation, Data curation, Writing - review & editing.

R.J.T: Funding acquisition, Investigation, Methodology, Resources, Writing – review & editing.

T.K: Funding acquisition, Investigation, Data curation, Writing - review & editing. M.P.M: Funding acquisition, Investigation, Data curation, Writing - review & editing.

A.S-A: Conceptualization, Data curation, Formal analysis, Funding acquisition, Investigation, Methodology, Project Administration, Resources, Visualization, Writing – original draft, Writing – review & editing.

B.D: Conceptualization, Data Curation, Formal analysis, Funding Acquisition, Project administration, Writing – review & editing.

## Supporting information

Supplementary Detailed Method 1

Supplementary Detailed Method 2

## Acknowledgments

Our thanks to the staff at SAHMRI LARIF/PIRL at Gilles Plains for their research support. Thank you to Dr. Adele Chow for her efforts in proofreading the final manuscript.

## Data Availability Statement

The underlying data supporting the findings of this study are available from the corresponding author upon reasonable request.

