## Supplementary Table 1 for "Optical Brain Pulse Monitoring for Detection of Large Vessel Occlusion Stroke in a Sheep Model"

**Supplementary Table 1.** *Arterial blood gas and systemic physiological parameters measured before and after induced stroke in Merino wethers.* Values are presented as median, minimum, and maximum for each parameter. Parameters include pH, partial pressure of oxygen ($pO_2$), partial pressure of carbon dioxide ($pCO_2$), bicarbonate ($HCO_3^-$), hemoglobin (Hb), hematocrit (Hct), sodium ($Na^+$), potassium ($K^+$), mean arterial blood pressure (MABP), pulse rate, and body temperature (oTemp). The data illustrate that arterial blood gas and systemic physiological parameters were maintained within normal limits throughout the experiment.

| | **Pre-Stroke** | | | **Post-Stroke** | | |
|---|---|---|---|---|---|---|
| | Median | min | max | Median | min | max |
| pH | 7.44 | 7.37 | 7.74 | 7.43 | 7.37 | 7.51 |
| $pO_2$ (mmHg) | 140.75 | 132.33 | 249.25 | 142.00 | 132.33 | 206.33 |
| $pCO_2$ (mmHg) | 42.00 | 37.00 | 47.00 | 42.00 | 37.00 | 47.00 |
| $HCO_3^-$ (mEq/L) | 28.08 | 24.63 | 31.40 | 27.90 | 23.55 | 31.40 |
| Hb | 8.00 | 5.00 | 9.70 | 8.00 | 5.00 | 9.70 |
| Hct (%) | 24.00 | 15.00 | 29.00 | 24.00 | 15.00 | 29.00 |
| $Na^+$ (mEq/L) | 140.50 | 136.50 | 147.00 | 140.00 | 135.00 | 145.00 |
| $K^+$(mEq/L) | 3.10 | 2.30 | 3.80 | 3.15 | 2.30 | 4.50 |
| MABP (mmHg) | 114.00 | 84.00 | 130.50 | 115.33 | 79.50 | 130.50 |
| Pulse (bpm) | 99.75 | 78.00 | 142.00 | 97.00 | 78.00 | 132.33 |
| oTemp (C°) | 37.40 | 33.85 | 39.30 | 37.30 | 33.85 | 38.23 |
