## Supplementary Detailed Method 1 for "Optical Brain Pulse Monitoring for Detection of Large Vessel Occlusion Stroke in a Sheep Model"

SUPPLEMENTAL MATERIALS

Supplementary Detailed Method 1. Surgical Approach

The ovine model of transient MCAo was implemented in this study as previously described[1,2]. To induce ischemic stroke, animals were positioned in the sphinx position with their heads tilted 90° for a right MCA approach. A 5 cm vertical incision was made, equidistant between the right eye and ear, terminating just below the zygomatic arch. The underlying branches of the superficial temporal artery were cauterized, and the large fat pad behind the orbit was removed to improve visualization.

The temporalis and other muscles of mastication were then stripped from the coronoid process of the mandible, and the coronoid was subsequently fractured at its base, adjacent to the zygomatic arch. The remaining muscles of mastication were stripped to expose the pterion. With the assistance of loupe magnification, a craniotomy was performed over the junction of the parietal and squamous temporal bones using a high-speed pneumatic drill (Midas Rex, Medtronic, USA), taking care not to breach the underlying dura. Bone was also removed anteroinferiorly from the greater wing of the sphenoid, and the craniotomy was extended extradurally with a Kerrison rongeur to provide access to the anterior temporal lobe beneath.

A horseshoe-shaped durotomy was then performed with an inferiorly based flap. Subpial MCA branches were identified, neurosurgical textile pads inserted over the exposed brain to prevent contusion, and gentle upwards retraction of the anterior temporal lobe was performed to follow each MCA branch to its bifurcation point. An aneurysm clip (Aesculap YASARGIL Aneurysm Clip, Germany) was then applied to the proximal MCA and left in situ for 4 hours. Synthetic dural regeneration matrix (Durepair, Medtronic, Australia) was used to cover the brain, autologous bone from the craniotomy reinserted, and overlying muscles opposed to enable $StO_2$% measurement throughout the MCAo period. To achieve reperfusion, the craniotomy site was re-opened, and the aneurysm clip was removed. Synthetic dura was then reapplied, a permanent cranioplasty was performed with autologous bone and polyacrylic cement (PMMA, Lang Dental, USA), and the overlying muscle was reopposed.
