## Supplementary Figure 1 for "Optical Brain Pulse Monitoring for Detection of Large Vessel Occlusion Stroke in a Sheep Model"

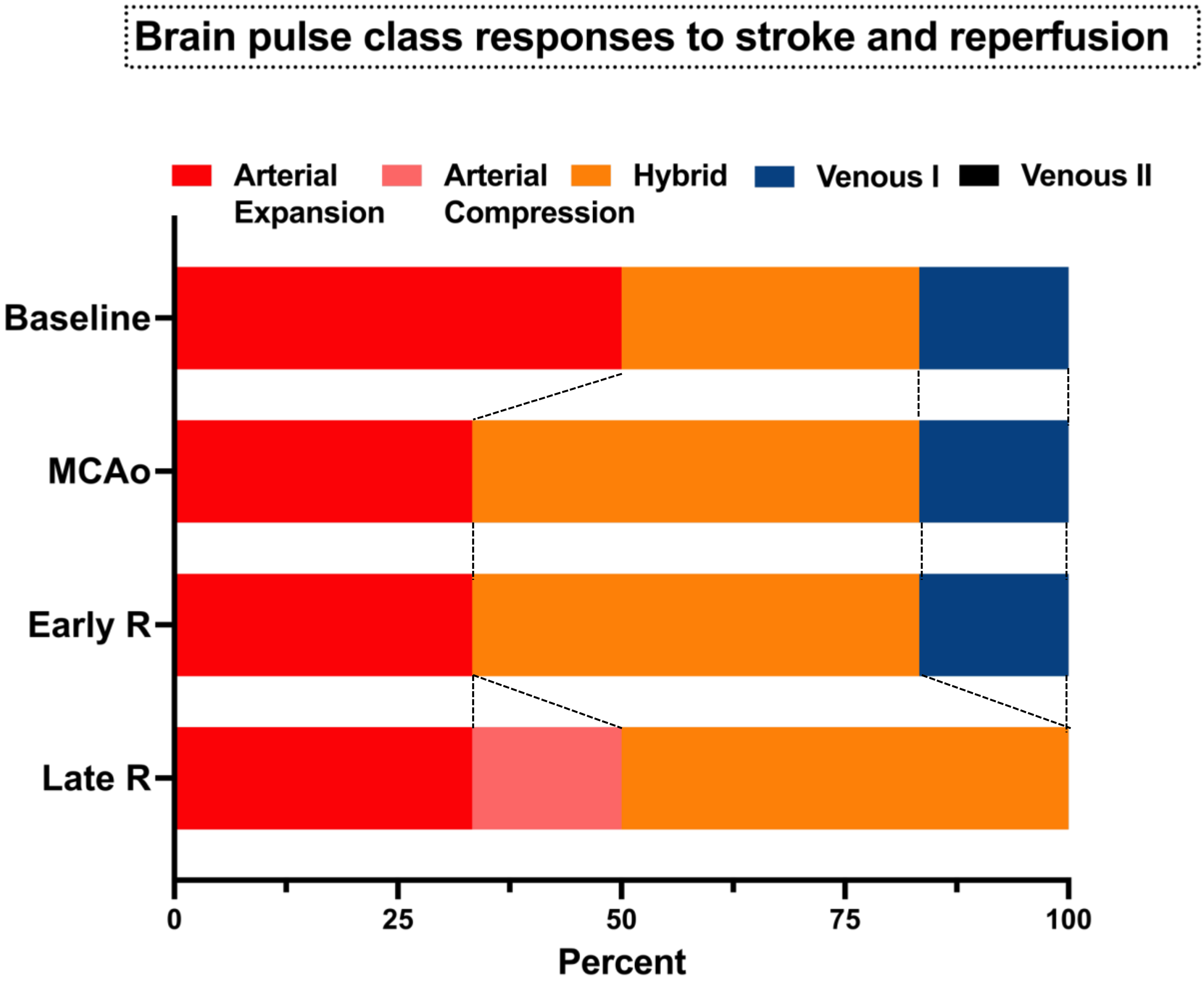


**Supplementary Figure 1. Contralateral hemisphere Brain pulse waveform classes responses to stroke and reperfusion.** The brain pulse classes did not change significantly during the study period. *Abbreviations: MCAo, middle cerebral artery occlusion ; ER, early reperfusion; LR, late reperfusion*.
