## Supplementary Detailed Method 2 for "Optical Brain Pulse Monitoring for Detection of Large Vessel Occlusion Stroke in a Sheep Model"

Supplementary Material 2. Detailed OBPM Signal processing

Data from the brain pulse monitor was captured at a rate of 500 Hz and stored for subsequent offline analysis. This analysis was conducted using Python v3.10, SciPy v1.10 [3,4], and various proprietary scripts within the Python environment. The primary objective was to ensure the integrity of signal analysis through an automated process that identifies noise characteristics based on an exhaustive array of statistical parameters.

The OBPM datasets were partitioned into discrete segments, each spanning 10 seconds, with a sliding window of 1 second to facilitate automated artifact detection and epoch rejection. The initial screening was conducted automatically, identifying epochs with raw signal amplitudes exceeding 2,000,000 units as saturated. Additionally, epochs were automatically flagged as compromised if they exhibited uniform values over more than 20 per cent of their duration, indicative of a flat signal profile.

Following the initial screening, the datasets were subjected to a high-pass filter using SciPy's Butterworth zero-phase filter (filtfilt) with a 1Hz cutoff and a 3rd order setting, aiming to eliminate signal drift. An automated noise detection algorithm then performed a comprehensive evaluation of the signal's statistical characteristics, including kurtosis, skewness, variance, mean, median, and their derivatives. These statistical measures were compared against predetermined thresholds to identify extreme anomalies suggestive of unacceptable noise levels. The Shapiro-Wilk test was also utilized automatically to assess the normality of the signal distribution, further enhancing the detection of noise-related irregularities.

The indices of automatically rejected epochs were used to excise corrupted signals from the raw dataset, which was then reassembled. To ensure continuity in the signal representation, segments showing significant amplitude discrepancies, as defined by a preset threshold, were automatically realigned starting from the terminal portion of the data. This realignment process was designed to preserve the integrity of the raw signal. To maintain the original dataset's length, the indices of the rejected epochs were reintegrated at their respective positions as NaNs (Not a Number). The preprocessing of signals was conducted independently across different channels, ensuring the procedure did not introduce cross-channel biases and maintained the data's fidelity. The preprocessed OBPM data was subsequently analyzed by specialized $StO_2$% prediction algorithms to generate oxygen saturation levels.
